# Increased utrophin expression in healthy and DMD patient derived myoblasts in response to ERK1/2 and EZH2 inhibitor treatment

**DOI:** 10.64898/2026.04.13.718206

**Authors:** Hannah J Gleneadie, Thomas Francis, Stephanie P L Mo, Aisha Ahmed, Mona Bensalah, Francesco Muntoni, Stephen D.R. Harridge, Matthias Merkenschlager, Amanda G Fisher

**Affiliations:** Epigenetic Memory Group, MRC Laboratory of Medical Sciences, Imperial College London, Hammersmith Hospital Campus, Du Cane Road, London, W12 0HS, UK; Centre for Human & Applied Physiological Sciences, Faculty of Life Science & Medicine, Kings College London, Shepherd’s House, Guy’s Campus, Great Maze Pond, London, SE1 1UL, UK; Roving Researcher, MRC Laboratory of Medical Sciences, Imperial College London, Hammersmith Hospital Campus, Du Cane Road, London, W12 0HS, UK; The Dubowitz Neuromuscular Centre, UCL Great Ormond Street Institute of Child Health, London, UK; MyoLine platform, Sorbonne Université, Inserm, Institut de Myologie, Centre de Recherche en Myologie, Paris, France; National Institute for Health Research Great Ormond Street Hospital Biomedical Research Centre, London, UK; Lymphocyte Development Group, MRC Laboratory of Medical Sciences, Imperial College London, Hammersmith Hospital Campus, Du Cane Road, London, W12 0HS, UK; Department of Biochemistry, University of Oxford, Oxford OX1 3QU, United Kingdom

**Keywords:** Duchenne Muscular Dystrophy, Dystrophin, Utrophin, Myoblasts, ERK1/2, EZH2, Primary human culture, Myopathy, inhibitor treatment, MyoD1

## Abstract

**Background:** The X-linked muscle wasting disorder Duchenne muscular dystrophy (DMD) is a progressive and ultimately fatal disease caused by loss of function mutations in the dystrophin (*DMD*) gene. Upregulation of utrophin (*UTRN*), an embryonic homologue of dystrophin, has been proposed as a therapeutic option that could ameliorate disease. We previously generated a bioluminescent screen for utrophin-upregulating compounds using a mouse reporter of endogenous utrophin expression and discovered that inhibition of ERK1/2 and EZH2, increases utrophin expression in myoblasts.

**Methodology:** Here we extend this analysis to show that treatment of human myoblasts with the ERK1/2 inhibitor LY3214996 and the EZH2 inhibitor GSK503, increases *UTRN* expression in primary and immortalised myoblasts derived from healthy volunteers and DMD patients.

**Results:** Short-term (24 hours) inhibition of ERK1/2 and EZH2 resulted in increased expression of utrophin in proliferating myoblasts. Surprisingly, in patient-derived samples, but not healthy controls, increased *UTRN* expression was sustained following drug removal and in vitro differentiation. Furthermore, dystrophin deficient myoblasts have altered expression of myogenic transcription factors *MYOD1* and *MYOG* and proliferation marker Ki67, signalling an altered regenerative capacity of these cells, while ERK1/2 inhibition, alone or combined with EZH2i, reversed this transcriptional signature.

**Conclusions:** Treatment with ERK1/2 and EZH2 inhibitors could offer a therapeutic option for DMD by increasing *UTRN* and *MYOD1* expression. We propose that this may compensate for *DMD* loss and help restore productive muscle differentiation and regeneration.

## Introduction

Muscle tissue is essential for movement, respiration and blood flow. The stimulus required for these processes is generated by repeated cycles of contraction and relaxation, resulting in considerable mechanical stress to the sarcolemma. To maintain integrity, the sarcolemma is connected to both the cytoskeleton and the extracellular matrix by multi-protein complexes such as the dystrophin-associated protein complex (DAPC) ^1^. The DAPC is organised around dystrophin. A cysteine-rich region at the C-terminal of dystrophin binds to the DAPC transmembrane protein β-dystroglycan, while the N-terminal binds to the actin cytoskeleton ^1-3^. In the absence of dystrophin, the DAPC dissociates and the connection between the extracellular matrix and the actin cytoskeleton is lost ^1^. In humans, this causes Duchenne muscular dystrophy (DMD), an X-linked muscle-wasting disorder caused by loss-of-function mutations in the dystrophin (*DMD*) gene ^4^. DMD is a progressive disease, patients begin to develop symptoms around the age of 2-3, become wheelchair bound around 10-12 and ultimately die at around 30 due to respiratory failure or cardiomyopathy ^5^.

In mice, loss of dystrophin (*Dmd*) results in a much milder phenotype ^6,7^. *Mdx* mice, which lack a full-length copy of the muscle-specific dystrophin protein ^8^, experience milder skeletal myopathy than DMD patients, late-onset or non-existent cardiomyopathy, and a near-normal lifespan ^6,7^. This is partially attributed to more effective muscle repair where an initial bout of degeneration in *mdx* mice is corrected by compensatory regeneration ^9^. Upregulation of utrophin (*UTRN, Utrn*), a homologue of dystrophin, has been proposed to be part of this protective mechanism. Accordingly, loss of both dystrophin and utrophin results in mice with a pathology that more closely phenocopies DMD ^7,10^. Utrophin is expressed at the sarcolemma during embryogenesis, however, as dystrophin expression increases, utrophin decreases and postnatally utrophin becomes confined to neuromuscular and myotendinous junctions (NMJ and MTJ respectively) ^11-13^. In *mdx* mice, utrophin can relocate to the sarcolemma and bind to components of the DAPC, partially compensating for dystrophin loss. This led to the hypothesis that utrophin upregulation could compensate for dystrophin loss in DMD patients. Accordingly, in *mdx* mice and a dystrophin-deficient canine model transgenic overexpression of utrophin prevents development of the dystrophic phenotype ^14-18^.

The search for small molecules to upregulate utrophin for the treatment of DMD is ongoing. Ezutromid, the most promising candidate identified to date, made it to clinical trials but failed to progress past stage II ^19,20^. The search is hindered by limitations in preclinical models and the complex regulation of utrophin expression. Multiple promoters and an upstream enhancer control *Utrn* / *UTRN* transcription ^21^, sites within the 3’UTR regulate mRNA stability and sites within the 5’ UTR regulate IRES-dependent translation ^22^. Previously, we described the generation of bioluminescent reporter mouse lines for dystrophin and utrophin, which track endogenous transcription and translation without impacting protein function ^23^. Reporter myoblasts were used in a pilot screen for utrophin-upregulating compounds, leading to the identification of EZH2 and ERK1/2 inhibition as a potential therapeutic option ^23^. Here, we use immortalised and primary myoblasts to examine the effect of ERK1/2 and EZH2 inhibitors on *UTRN* expression in human cells. We validate the efficacy of the screening platform previously described by showing that ERK1/2 and EZH2 inhibitor treatment can increase *UTRN* expression in human myoblasts. We found comparable changes to *UTRN* expression in proliferating myoblasts derived from healthy volunteers and DMD patients. However, following drug removal and in vitro myogenesis, *UTRN* upregulation is lost in healthy samples but maintained in DMD myoblasts. Moreover, loss of dystrophin in human myoblasts alters the expression of differentiation and proliferation markers, while ERK1/2 inhibition, particularly when combined with EZH2 inhibition, can restore these. Therefore, ERK1/2 and EZH2 inhibition could be a potential therapeutic option for DMD.

## Results

Previouslys we have shown that treatment with ERK1/2 inhibitor LY3214996 (LY32) and EZH2 inhibitor GSK503, alone or more potently combined, cause increased *Utrn* expression in mouse myoblasts ^23^. To ask whether enhanced *UTRN* expression was also evident in human cells, we treated human myoblasts with LY32 and GSK503. Primary human myoblasts were isolated from 6 healthy young adult males (aged 20-32 years) and proliferating myoblasts were then treated with increasing concentrations of LY32 and GSK503 for 24 hours. *UTRN* mRNA showed a quantitative dose-dependent response to treatment, with particularly increased levels when both drugs were applied at the highest concentration tested (10 μM) (Figure 1 a). All results were normalised to TATA box binding protein (TBP), which is stably expressed in myoblasts ^24^ and *UTRN* mRNA levels are shown relative to vehicle only-samples.

**Figure 1:**
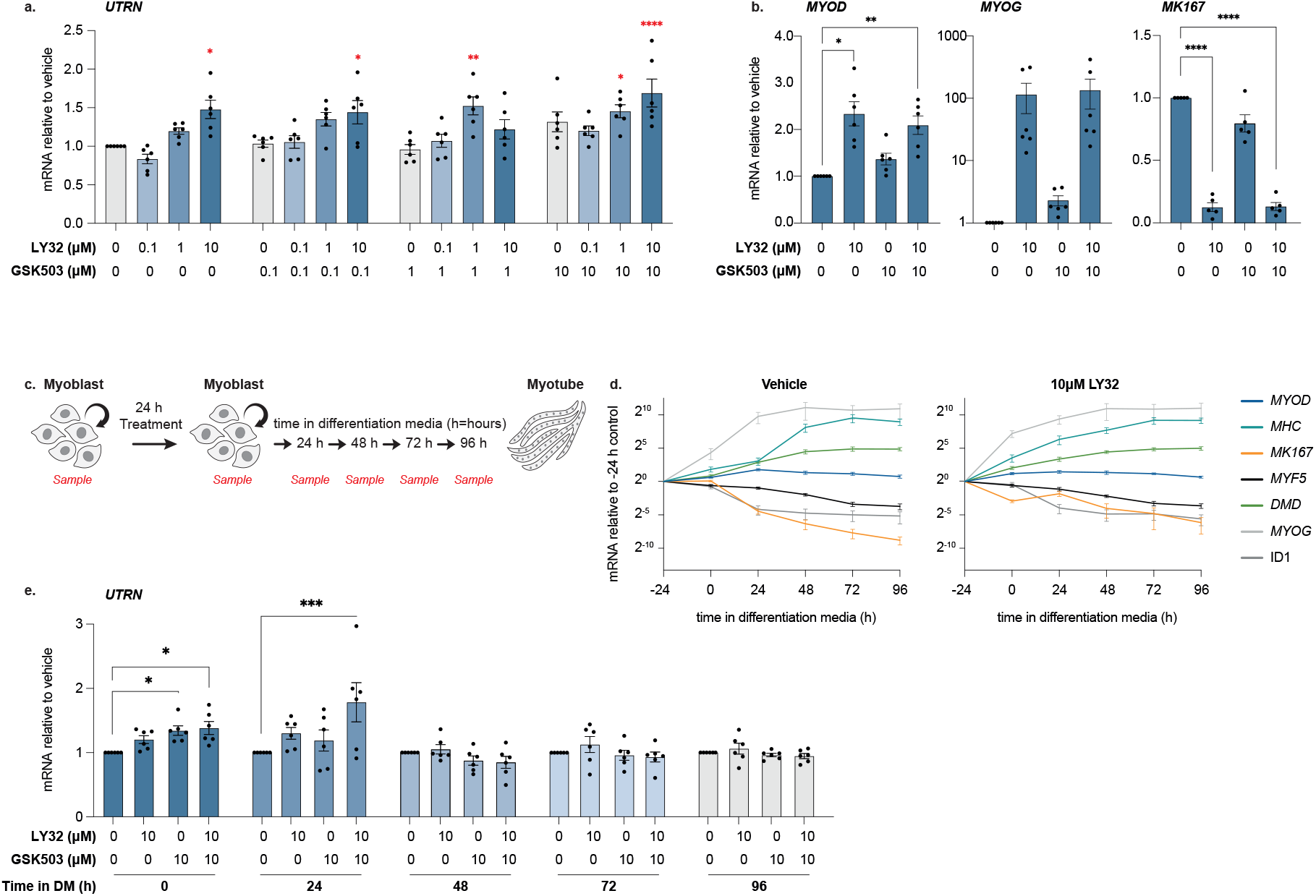
Impact of ERK1/2 and EZH2 inhibitor treatment on gene expression in healthy human myoblasts. Primary human myoblasts isolated from healthy young adult males (aged 20-32 years) were treated with ERK inhibitor LY3214996 (LY32) and EZH2 inhibitor GSK503 for 24 hours. n=6 independent biopsies. **a**. RT-qPCR of *UTRN* following titrations of LY32 +/- GSK503. **b**. RT-qPCR for *MYOD1, MYOG* and *MK167* following 10 μM treatment with GSK503 +/- LY32 or vehicle. **a-b**. Results are shown relative to the vehicle-treated sample. One-way ANOVA with Šidák’s multiple comparison test compares vehicle to inhibitor treated samples. **c**. Schematic representing the treatment and sampling scheme used in **d**,**e** and **supplementary figure 1**. Proliferating myoblasts were sampled prior to treatment, exposed to inhibitors for 24 hours and sampled again. Inhibitors were removed and myoblasts transferred to differentiation media (DM). Myoblasts were differentiated into myotubes for 96 hours in differentiation media. Samples were taken every 24 hours. **d**. mRNA was extracted from each timepoint and RT-qPCR for markers of myogenic differentiation performed. Results are shown relative to the untreated, undifferentiated samples (-24 h in DM). Graph is in Log2 scale. **e**. RT-qPCR for *UTRN* mRNA for samples collected as of **c**. For each time point, treated samples are shown relative to the untreated sample of that time point. A 2-way ANOVA with Dunnett’s multiple comparison test compares vehicle with treated samples matched to each time point. **a**,**b**,**d**,**e**: Results are normalised to *TBP*. All graphs show mean (n=6) +/- SEM. *p<0.05, **p<0.01, ***p<0.001, ****p<0.0001.

We, and others, have shown that ERK inhibition can cause premature expression of myoblast differentiation markers ^23,25^. This result was replicated in proliferating human myoblasts; 24-hour treatment with LY32 increased expression of myogenic transcription factors *MYOD1* and *MYOG* (myogenin) and decreased expression of proliferation marker *MK167* (Ki67), while inhibition of EZH2 had minimal effects (Figure 1 b). Interestingly, when treatment was removed and the cells allowed to differentiate in low serum media, both treated and untreated myoblasts formed myotubes. Moreover, when pre-treated cells were sampled throughout myogenesis (as outlined in Figure 1 c), myogenic markers were similarly expressed in LY32 pre-treated and untreated cells following 2 days culture in differentiation medium (DM) (Figure 1 d), suggesting that while ERK inhibition promotes premature expression of differentiation markers, it ultimately does not interfere with in vitro myogenesis. Indeed, when *MYOD1* and *MYOG* mRNA levels were normalised to time-matched vehicle-treated controls, there was no difference in expression between LY32-treated and vehicle-treated samples following 24 h culture in DM (Supplementary Figure 1 a, left and middle panel). Similarly, the percentage of myogenin positive nuclei, as determined by immunofluorescence staining, was unaltered after 24 h in DM (Supplementary Figure 1 b). Proliferation, represented by *MK167* mRNA (Supplementary Figure 1 a, right) and immunofluorescence labelling of Ki67 (Supplementary Figure 1 c, quantified in d) decreased following LY32 treatment. However, 24 hours after drug and serum withdrawal, *MK167* mRNA and Ki67 nuclear intensity were higher in pre-treated myoblasts, suggesting increased proliferation in LY32-pretreated cells after the onset of differentiation.

Dystrophin performs essential functions in both differentiated myofibres and myogenic progenitors ^26,27^. Hence, a successful treatment option for DMD would ideally target both cell types. We have previously shown in healthy mouse myoblasts that proliferating cells were susceptible to LY32 and GSK503 treatment, while differentiated cells were not. However, mouse cells pre-treated with LY32 and GSK503 maintained increased *Utrn* mRNA following drug withdrawal and differentiation ^23^. Here, we show that LY32 and GSK503 treatment can increase *UTRN* expression in proliferating human primary myoblasts, however, this is no longer apparent after 48 hours of differentiation (Figure 1 e).

To address whether *UTRN* upregulation in response to ERK1/2 and EZH2 inhibition is also induced in DMD patient-derived myoblasts we obtained immortalised myoblasts (from males aged 13-16 years, details in Supplementary Figure 2 a) from the Centre de Researche en Myologie (Paris). Consistent with previous results, healthy immortalised myoblasts (AB1190) upregulated *UTRN* mRNA in response to ERK1/2 and EZH2 inhibition (Figure 2 a, blue bars). *UTRN* mRNA was also upregulated in DMD myoblast line AB1098 and validated in a further DMD line, AB1071 (Figure 2 a, orange bars). Utrophin protein, as determined by western blot analysis, also increased upon treatment with LY32 and GSK503 in both AB1190 and AB1098 cells (Figure 2 c, and quantified in b). As expected, dystrophin was not observed in DMD myoblasts (AB1098) (Figure 2 c, and quantified in b). However, in the healthy control line (AB1190) dystrophin was upregulated upon LY32 treatment (Figure 2 c, and quantified in b), likely due to precocious expression of myogenic transcription factors following ERK1/2 inhibition. To validate these results, we obtained 16 primary DMD patient-derived myoblast cultures from the MRC Centre for Neuromuscular Diseases (MRC CNMD) Biobank (London). Donors ranged in age from 1 to 12 years and had multiple exon deletions in the *DMD* gene (between exons 42-54) (details in Supplementary Figure 2 b). Consistent with the cell line experiments, all primary myoblasts from the DMD patients upregulated *UTRN* mRNA in response to LY32 and GSK503 treatment; with *UTRN* increased over 1.5-fold in 10 samples following LY32 treatment and over 2-fold in 6 samples following combination therapy. When the response is viewed as a population average, 24-hour treatment with LY32 and GSK503 significantly increased *UTRN* mRNA expression in primary DMD myoblasts (Figure 2 d). This was accompanied by a corresponding increase in *MYOD1* and *MYOG* and a decrease in *MK167* mRNA (Figure 2 e). Interestingly, for LY32 and combined treatment, there was a slight trend towards higher *UTRN* upregulation in samples derived from older patients (Figure 2 f), while the size and location of the disease-causing mutation did not have a discernible effect.

**Figure 2:**
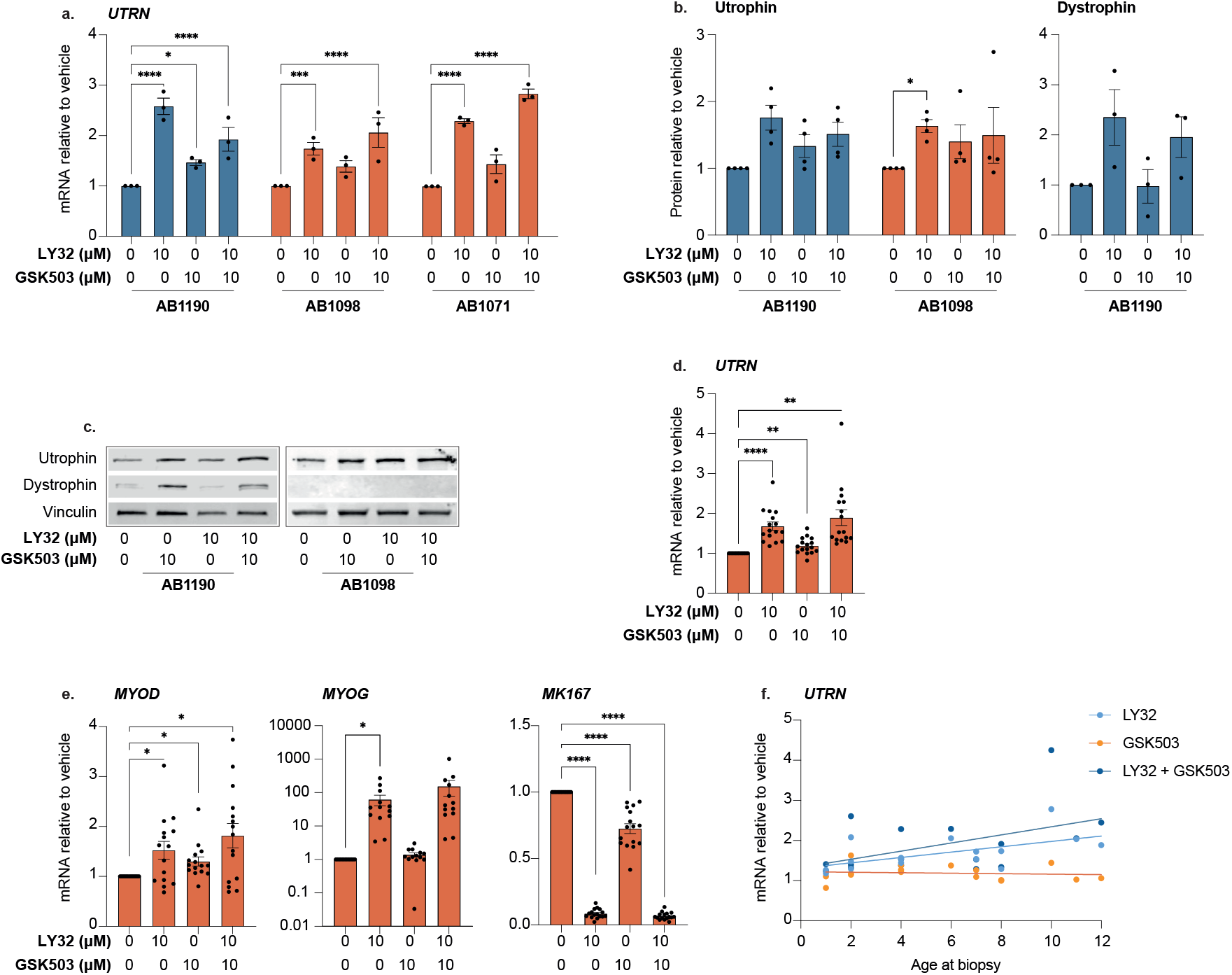
*UTRN* upregulation in DMD patient-derived myoblasts following LY32 and GSK503 treatment. **a-c**. Immortalised myoblasts from DMD patients (orange bars, AB1098 and AB1071) and healthy control (blue bars, AB1190) were treated with 10 μM LY32 +/- 10 μM GSK503 or vehicle (DMSO) for 24 hours. For each cell line and condition, n=3. **a**. RT-qPCR for *UTRN* mRNA following treatment. *UTRN* is normalised against *TBP* and samples are shown relative to the vehicle-only control. A two-way ANOVA with Šidák’s multiple comparison test was used to compare vehicle with treated samples. **b-c**. Western blot analysis of utrophin and dystrophin protein following treatment. Representative images (**c**) and quantification of signal (**b**). Utrophin and dystrophin protein levels are normalised to the corresponding loading control, vinculin and relative to the vehicle only control. **d-f**. Primary myoblasts isolated from DMD patients were treated with 10 μM LY32 +/- 10 μM GSK503 or vehicle (DMSO) for 24 hours. n=16 independent biopsies. RT-qPCR for *UTRN* (**d**) and *MYOD1, MYOG* and *MK167* (**e**) mRNA following treatment. Results are normalised against *TBP* and shown relative to the vehicle treated sample. One-way ANOVA with Dunnett’s multiple comparison test compares vehicle with treated samples. **f**. Results from **d**. are shown relative to the age of the patient at the time of biopsy. **p<0.01, ***p<0.001, ****p<0.0001. All graphs represent the mean +/- SEM.

Dystrophin deficient myoblasts are phenotypically distinct from healthy cells, with altered myogenic and proliferative potential ^28^. Accordingly, we observe reduced mRNA levels of myogenic transcription factors *MYOD1* and *MYOG* in DMD myoblast lines, AB1098 and AB1071 (orange), compared to the healthy control line, AB1190 (blue) (Figure 3 a and b, respectively) and an equivalent trend of lower *MYOD1* in primary DMD myoblasts compared to healthy controls (Supplementary Figure 3 a). Conversely, immunofluorescent staining of Ki67, a marker of proliferation, was higher in the DMD-derived line (representative images in Figure 3 c, and quantified in d). Encouragingly, treatment with LY32, particularly when combined with GSK503, can restore some of these phenotypic differences. In DMD myoblasts AB1098 treatment leads to increased *MYOD1* and *MYOG* mRNA (Figures 3 e and f, respectively) while Ki67 nuclear intensity was significantly decreased (Figure 3 g, quantified in h). Altered proliferation is confirmed by a decrease in cell number following treatment that is maintained 24 hours after drug withdrawal but returns to near vehicle-treated levels following one week in culture (Supplementary Figure 3 b). Utrophin has been shown to increase during in vitro myogenesis ^21,23^, therefore the increase in *UTRN* following LY32 and GSK503 treatment may reflect the premature expression of myogenic differentiation markers. Knockdown of JAK1 is known to promote premature myogenic differentiation ^29^. Accordingly, treatment with ruxolitinib (Ruxo), a JAK1/2 inhibitor, increased *MYOD1* mRNA in both AB1098 and AB1190 myoblasts (Supplementary Figure 3 c) and increased *UTRN* mRNA in DMD patient-derived myoblasts but not healthy controls (Figure 3 i).

**Figure 3:**
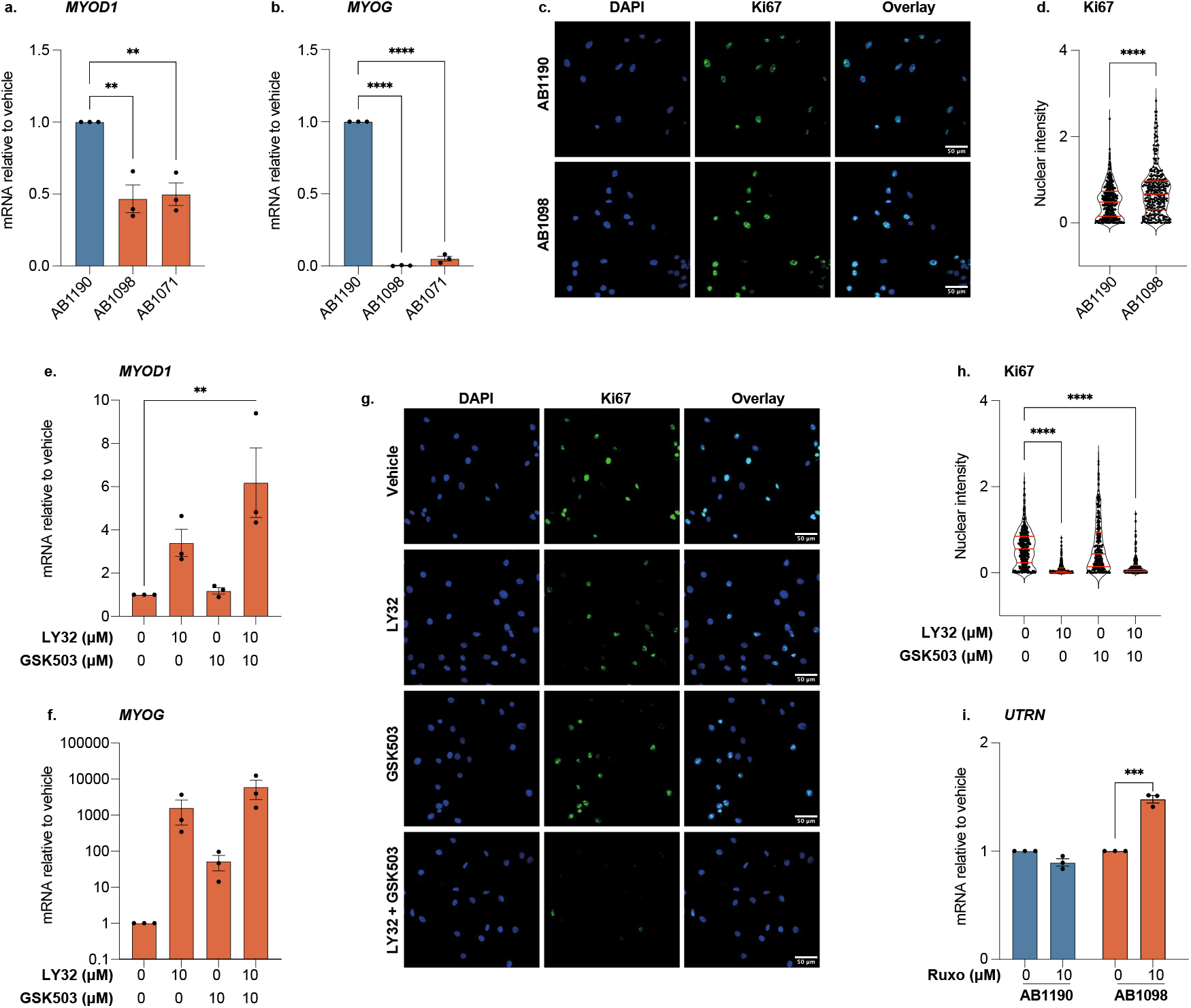
Markers of proliferation and myogenesis are altered in DMD patient-derived myoblasts but restored following treatment with LY32 and GSK503. **a-b**. RT-qPCR of levels of *MYOD1* (**a**) and *MYOG* (**b**) in immortalised DMD myoblast cell lines AB1098 and AB1071 compared to healthy control, AB1190. One way ANOVA with Dunnett’s multiple comparison test was used to compare AB1190 to DMD cell lines. **c-d**. Immunofluorescent staining for Ki67 in AB1190 (upper) and AB1098 (lower) cells. **c**. representative images. **b**. Quantification of **c**. FIJI was used to calculate Ki67 intensity in nuclei. For each nucleus Ki67 signal was normalised against the corresponding DAPI signal. Quartiles and median lines are shown in red. An unpaired T-test was used to compare AB1190 to AB1098. **e-h**. AB1098 cells were treated with 10 μM LY32 +/- 10 μM GSK503 or vehicle for 24 hours. **e-f**. RT-qPCR of *MYOD1* (**e**) and *MYOG* (**f**) (shown in log10 scale) following inhibitor treatment. One way ANOVA with Dunnett’s multiple comparison test compares vehicle with treated samples. **g-h**. Immunofluorescent staining for Ki67 in AB1098 cells. **g**. Representative image. **h**. Quantification of **g** (as of **c**). **i**. AB1190 and AB1098 cells were treated with JAK1/2 inhibitor Ruxolitinib (Ruxo) for 24 hours. RT-qPCR for *UTRN* mRNA following treatment. Two-way ANOVA with Šidák’s multiple comparison test compares vehicle with treated samples. **a**,**b**,**e**,**f**,**i**: Results are normalised to *TBP* and shown relative to control line AB1190 (**a**,**b**) or vehicle-treated sample (**e**,**f**,**i**). Graphs show mean (n=3) +/- SEM. **p<0.01, ***p<0.001, ****p<0.0001.

Here, we have discussed how a number of differences between DMD and control myoblasts can be restored by LY32 and GSK503 treatment, therefore we sought to determine the impact of dystrophin deficiency on *UTRN* upregulation. DMD patients are known to upregulate utrophin to compensate for dystrophin loss ^30,31^. Indeed, AB1098 myoblasts have significantly higher *UTRN* mRNA than AB1190 cells (Figure 4 a) and primary DMD myoblasts tend towards higher *UTRN* expression than healthy controls (Figure 4 b). Moreover, distinct from healthy human myoblasts, when immortalised myoblasts were pre-treated with LY32 and/or GSK503 and sampled following drug removal and differentiation (outline of the treatment and sampling strategy in Figure 4 c), DMD myoblasts uniquely responded to treatment. Upregulation of utrophin mRNA (Figure 4 d) and protein (Figure 4 e, quantified in f) was sustained in DMD myoblasts AB1098, but not control line, AB1190, with an equivalent result in additional DMD cell line, AB1071 (Figure 4 d). This was further validated in primary culture. Pre-treatment of primary DMD myoblasts caused a sustained increase in *UTRN* mRNA in 11 of the 14 biopsies tested. When analysed as a population, LY32 and GSK503 pre-treatment resulted in significantly higher *UTRN* mRNA than vehicle pre-treated cells (Figure 4 g), contrary to the result for healthy myoblasts previously described (Figure 1 e). Therefore, like healthy mouse myoblasts ^23^, DMD patient-derived myoblasts are able to retain utrophin upregulation following drug withdrawal and in vitro myogenesis, while healthy human myoblasts are not (summarised in Supplementary Figure 4 a). The expression of myogenic markers *MYOD1* and *MYOG* did not differ between vehicle pre-treated and drug pre-treated myotubes for any of the cells tested (Supplementary Figure 4 b and c) and the increase in *UTRN* mRNA in AB1098 cells following ruxolitinib treatment was lost following drug withdrawal and differentiation (Figure 4 h). These results suggests that the discrepancy in the pre-treatment response between healthy and DMD patient derived myoblasts does not reflect differences in in vitro myogenesis and instead is likely linked to higher basal levels of *UTRN* mRNA in myoblasts derived from DMD patients.

**Figure 4:**
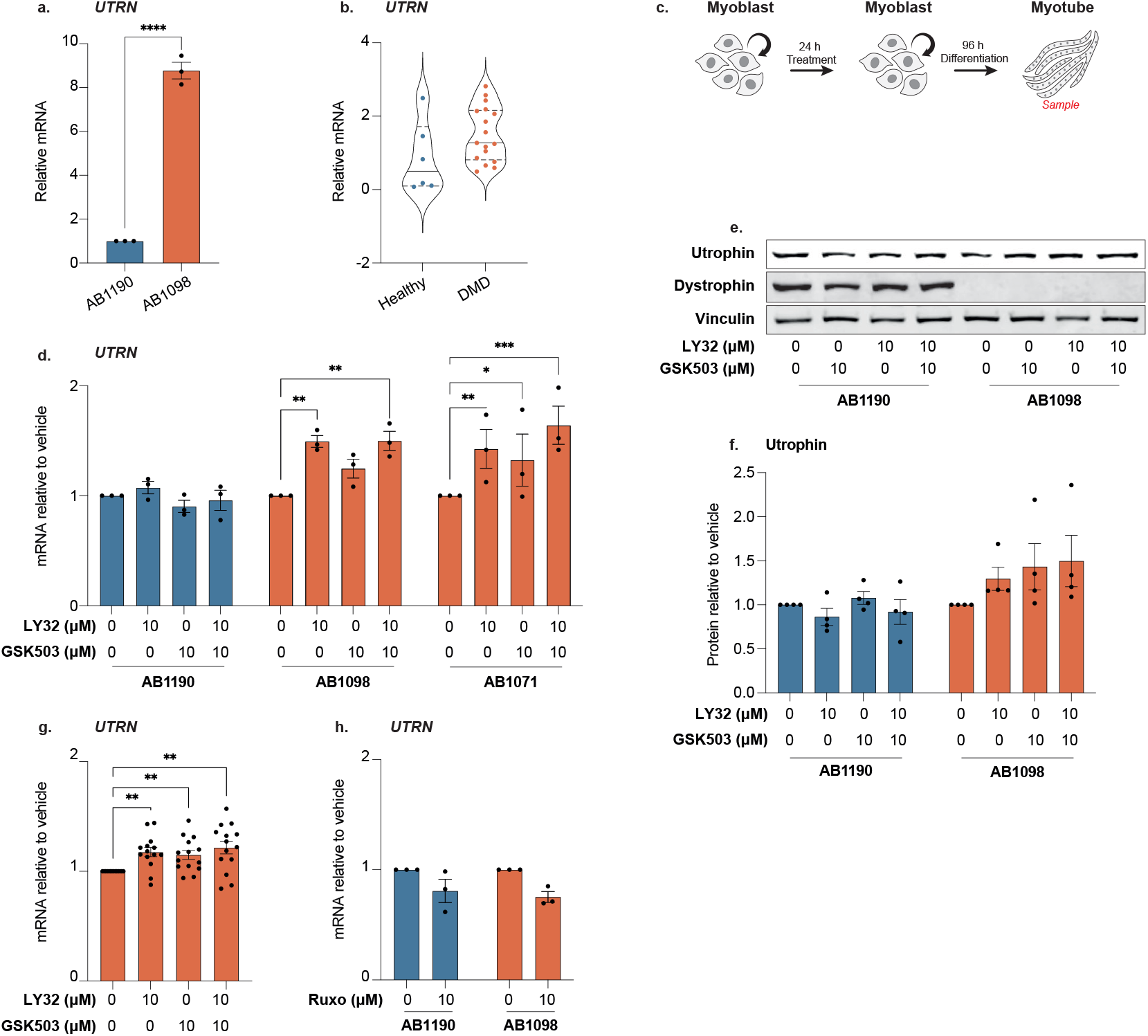
Pre-treatment of DMD patient-derived myoblasts with ERK1/2 and EZH2 inhibitors results in *UTRN* upregulation in resulting myotubes. **a,b**. RT-qPCR of basal levels of *UTRN* mRNA expression in healthy vs DMD patient-derived immortalised (**a**) (n=3 for each cell line) and primary (**b**) myoblasts. **b**. n=6 healthy myoblasts and n=16 DMD myoblasts. Lines show median and quartiles. **c**. Schematic representing the treatment and sampling strategy employed in **d-h** and **Supplementary Figure 4**. Proliferating myoblasts were treated with inhibitors for 24 hours, following this, treatment was removed and replaced with differentiation medium (DM). Cells were allowed to differentiate for 96 hours and sampled for RNA or protein. **d-f**. 10 μM LY32 +/- 10 μM GSK503 or vehicle pre-treatment in immortalised DMD patient-derived (orange bars, AB1098 and AB1071) and healthy control (blue bars, AB1190) myoblasts. **d**. RT-qPCR for *UTRN* mRNA following pre-treatment (n=3). Two-way ANOVA with Šidák’s multiple comparison test was used to compare vehicle with treated samples. **e-f**. Western blot, representative image of utrophin, dystrophin and vinculin protein (**e**) and analysis of utrophin levels (**f**). Utrophin protein is normalised against vinculin and shown relative to the vehicle control. **g**. RT-qPCR of *UTRN* mRNA in primary DMD patient-derived myoblasts (n=14) following 10 μM LY32 +/- 10 μM GSK503 or DMSO (vehicle) pre-treatment. **h**. Immortalised myoblasts AB1190 and AB1098 were pre-treated with Ruxolitinib for 24 hours prior to differentiation. RT-qPCR for *UTRN* mRNA following differentiation. **a**,**b**,**d**,**g**,**h**. Results are normalised to *TBP* and shown relative to vehicle pre-treated sample (**b, g, h**) or healthy controls (**a**). Graphs show mean +/- SEM. *p<0.05, **p<0.01, ***p<0.001.

## Discussion

Postnatal upregulation of utrophin in muscle tissue is thought to compensate for the lack of dystrophin in DMD patients, however, no compound or treatment regime has been found that translates this hypothesis to the clinic ^32^. In our previous work, we described the generation of a bioluminescent screening platform to identify compounds which upregulate utrophin in adult mouse myoblasts and determined that inhibition of ERK1/2 and EZH2 could increase *Utrn* ^23^. Here, we extend this observation to humans and show that ERK1/2 and EZH2 inhibition increases utrophin expression in human myoblasts isolated from both healthy volunteers and DMD patients.

In vivo, muscle regeneration requires myoblasts to proliferate, migrate to the source of damage, and differentiate. These three processes are altered in dystrophin-deficient myoblasts. Cells are found to have increased proliferative potential, disrupted chemotaxis and an altered readiness to differentiate ^28^. Here, we show that the myogenic transcription factors *MYOD1* and *MYOG* are decreased and the proliferation marker Ki67 increased in DMD myoblasts as compared to healthy controls. MyoD1 is a key myogenic transcription factor, essential for muscle differentiation, known to regulate myogenin expression and inhibit proliferation. Satellite cells lacking *Myod1* fail to differentiate ^33^, and in *mdx x Myod1*^*-/-*^ mice impaired regeneration exacerbates the dystrophic phenotype ^34,35^. In dystrophin-deficient myoblasts, reduced expression of *MYOD1* therefore probably reflects a compromised regenerative capacity. Combined with satellite cell division abnormalities that impact myoblast generation ^26^, poor regeneration will drive DMD progression, as dystrophic myoblasts fail to repair muscle damaged by myofibre instability ^28^. Here, we show that treatment with LY32, in particular when combined with GSK503, reverses some of the effects of dystrophin loss in myoblasts. Expression of *MYOD1* and *MYOG* are increased following treatment, while proliferation, indicated by Ki67 expression, is reduced. Corresponding upregulation of utrophin could be linked to this precocious expression of muscle differentiation markers. Accordingly, inhibition of JAK1/2, also thought to promote premature differentiation, lead to a similar increase in *UTRN* in DMD myoblasts ^29^.

The progressive loss of healthy muscle tissue in DMD patients has been largely attributed to the fragility of dystrophin-deficient myofibres and the accumulation of damage ^27^. However, dystrophin loss also affects the biology of muscle stem cells; lack of polarity causes deficits in myogenic progenitors ^26^ and repeated cycles of regeneration leads to depletion of the satellite cell pool ^36^. Interestingly, in absence of stressors such as eccentric exercise, loss of dystrophin in fully differentiated myofibres does not cause overt degeneration ^37^. Treatments which primarily target myogenic progenitors could therefore have a therapeutic effect, if functional myoblasts are able to differentiate into functional myofibres. This could convey a long-term cure for DMD if therapy were to be applied before the onset of pathology, before the secondary effects of myofibre necrosis and regeneration take hold ^37^. Previous results showed that LY32 and GSK503 treatment was ineffective at utrophin upregulation in differentiated mouse cells (cultured myotubes and ex vivo myofibres) ^23^. However, pre-treatment of proliferating mouse myoblasts with inhibitors prior to differentiation led to increased *Utrn* expression in the resulting myotubes ^23^. Intriguingly, we observed a similar sustained increase in utrophin expression in DMD patient-derived myoblasts, but not in healthy control cells. We hypothesise that this may reflect differences in basal utrophin levels between different cell types and consistent with this, we observe a trend towards higher utrophin in DMD patient-derived myoblasts than healthy cells. Further work into the chromatin landscape at the *UTRN* locus in healthy and dystrophic myoblasts will be required to establish the mechanistic basis of such differences.

The prior generation of a bioluminescent screening platform that dynamically reports endogenous utrophin transcription and translation, without impacting protein function represents a powerful new tool to identify utrophin upregulating compounds ^23^. Here we validated its efficacy by confirming that identified compounds also upregulate utrophin expression in healthy and DMD patient-derived human myoblasts. While, the discrepancy between mouse and human cells cautions against relying solely on mouse models for DMD research, our data encourages the iterative application of both models to progress effective treatments towards eventual human trials ^38^.

## Conclusions

Here, we have shown that LY32 and GSK503 treatment upregulates utrophin in healthy and DMD patient-derived human myoblasts. This is accompanied by an increase in myogenic transcription factors, and a decrease in Ki67, reversing some of the phenotypes associated with dystrophin deficiency. Moreover, in DMD myoblasts, but not healthy controls, utrophin upregulation is maintained following drug removal and in vitro myogenesis. Combined, these results are likely to be of therapeutic benefit for Duchenne muscular dystrophy patients: increased utrophin expression may compensate for dystrophin loss in myoblasts and resulting myofibres, while increased *MYOD1* may help restore the regenerative capacity of dystrophic myoblasts.

## Acknowledgements

This work was funded by the Medical Research Council (MRC) (A.G.F. by MC_PC_23024 and MC_UP_1605/12). T.F and S.D.R.H are supported by the Ageing Biology Foundation EU.

We would like to thank the Myoline Platform, from the Institut de Myologie, Paris, for the generous donation of the immortalised human myoblast lines. The MRC Centre for Neuromuscular Diseases Biobank London, supported by the National Institute for Health Research Biomedical Research Centres at Great Ormond Street Hospital for Children NHS Foundation Trust and at University College London Hospitals NHS Foundation Trust and University College London, is acknowledged for providing primary DMD myoblast cultures.

The authors would like to thank Holly Johnston and Dr Jamie Meredith (MRC Laboratory of Medical Sciences, London) for their help and support in obtaining myoblast samples. Finally, we would like to thank the Roving Researcher scheme at the MRC Laboratory of Medical Sciences (London) for providing an additional researcher (S.P.L. M) to support the project whilst H.J.G was on maternity leave.

## Author contributions

H.J.G and A.G.F conceptualised the study with help from T.F and S.D.R.H. Experiments were performed by H.J.G with help from T.F and S.P.L.M. S.P.L.M optimised western blot protocols and T.F helped with primary myoblast culture. T.F and S.D.R.H isolated primary myoblasts from healthy human volunteers. Immortalised myoblasts were generated by M.B. Primary DMD patient-derived myoblasts were provided by F.M and A.A. H.J.G and A.G.F wrote the manuscript with input from co-authors.

## Competing interests

All authors declare no competing interests.

## Data availability statement

Source data is provided in Supplementary Data file 1.

**Supplementary Figure 1:**
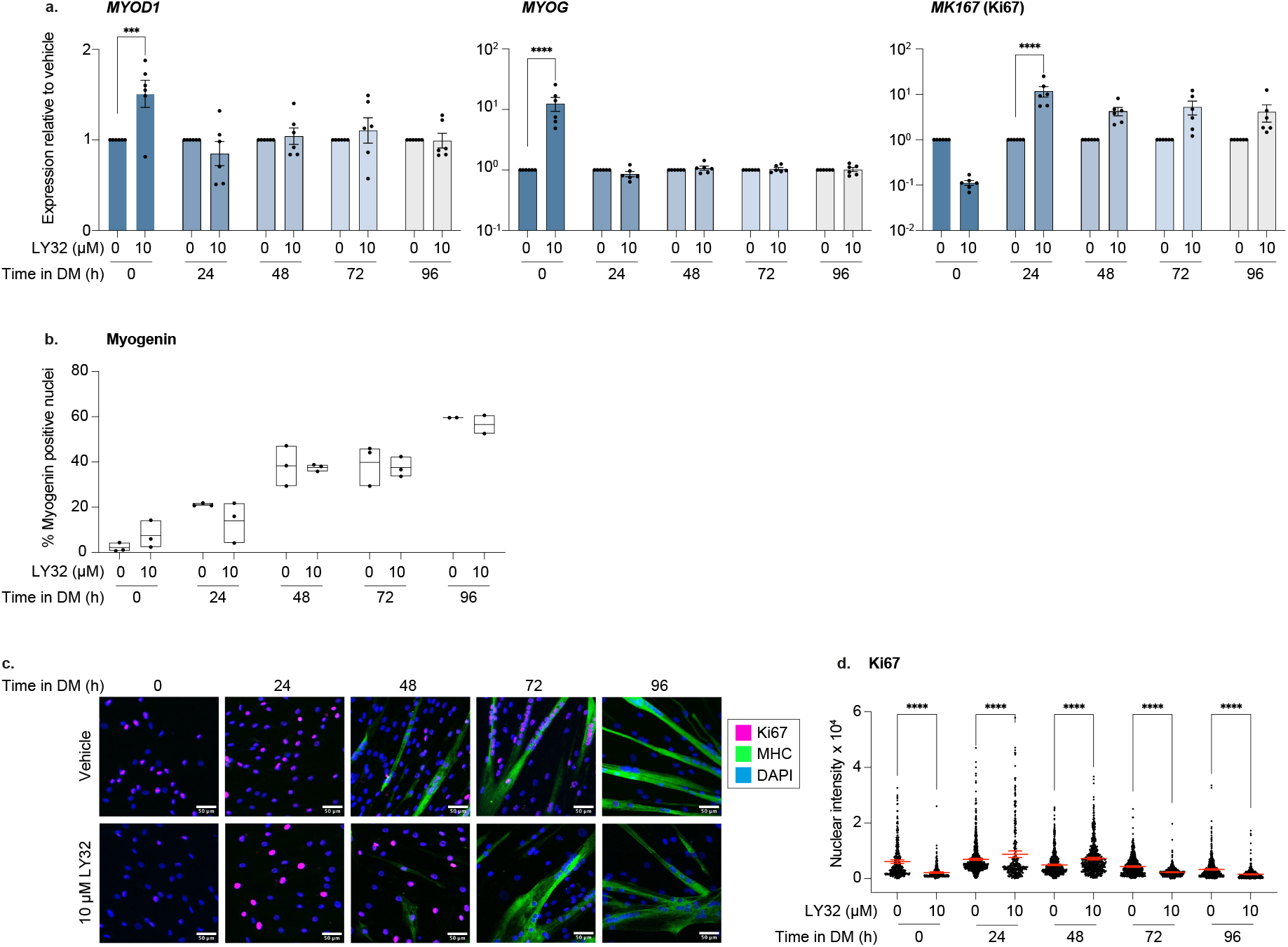
Human myoblasts are able to differentiate into myotubes following pre-treatment with ERK inhibitor LY32. **Corresponding to Figure 2**. following which media Primary human myoblasts were treated with 10 μM LY32 or vehicle for 24 hours, was removed and replaced with differentiation medium (DM). Cells were treated and sampled as outlined in **Figure 1c. a**. RT-qPCR for *MYOD1* (left), *MYOG* (middle) and *MK167* (right) mRNA. **b**. Quantification of myogenin immunofluorescent staining across differentiation. The number of myogenin positive nuclei is shown relative to the total number of DAPI positive nuclei. Results show the average of 3 independent biopsies, with 2 images analysed per biopsy. **c**. Representative images of immunofluorescent staining for Ki67 (Pink), MHC (Green) and DAPI (Blue). **e**. Quantification of Ki67 nuclear intensity in **c**. FIJI was used to assign nuclei based on DAPI staining and determine Ki67 intensity in each nucleus. Over 100 nuclei were analysed per condition. One-way ANOVA with Šidák’s multiple comparison test was used to compare LY32 pre-treated with vehicle pre-treated samples for each time point. **a, d**. Samples are normalised against *TBP* and for each time point, treated samples are shown relative to the time-matched vehicle treated control. A 2-way ANOVA with Dunnett’s multiple comparison test compares vehicle with treated samples matched to each time point. Graphs show mean (n=6) +/- SEM. ***p<0.001, ****p<0.0001.

**Supplementary Figure 2:**
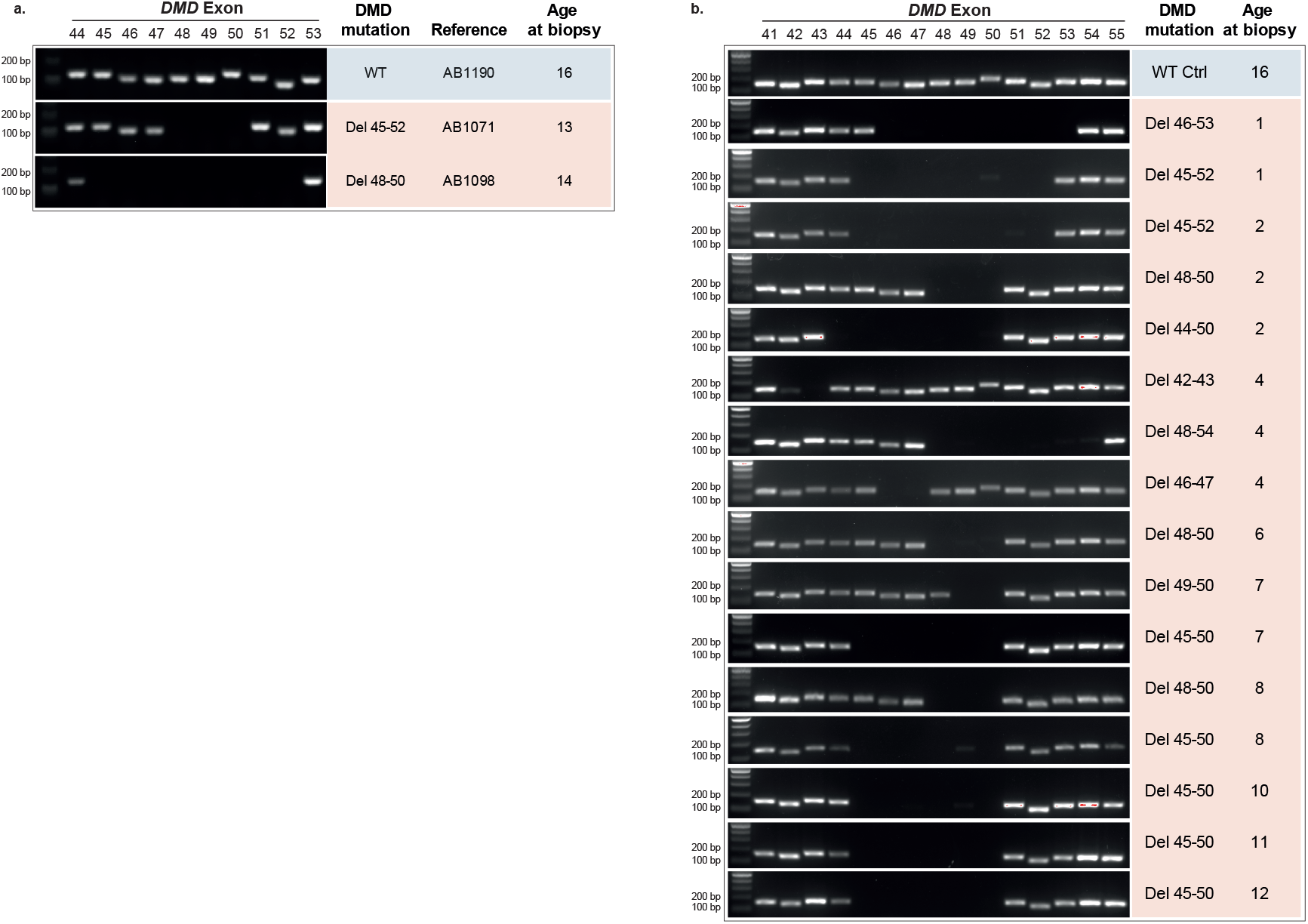
Genotyping of DMD patient-derived myoblasts. **Corresponding to Figure 2. a**. Genotyping for immortalised healthy (WT) (blue) and DMD patient-derived myoblasts (orange) from the Centre de Researche en Myologie. DNA was isolated and genomic PCR performed for DMD exon 44-53. The age of the patient at the time of biopsy, and the nomenclature used to refer to each line is shown alongside the genotyping result. **b**. DMD patient-derived myoblasts from the CNMD biobank contain multiple exon deletions in *DMD* between E41 to E55. Genotyping was performed as of **a**. Deletions correspond with that expected from patient data. Immortalised WT control line AB1190 is shown at the top (in blue) as a positive control for each PCR. PCR products all range between 100-200 bp.

**Supplementary Figure 3:**
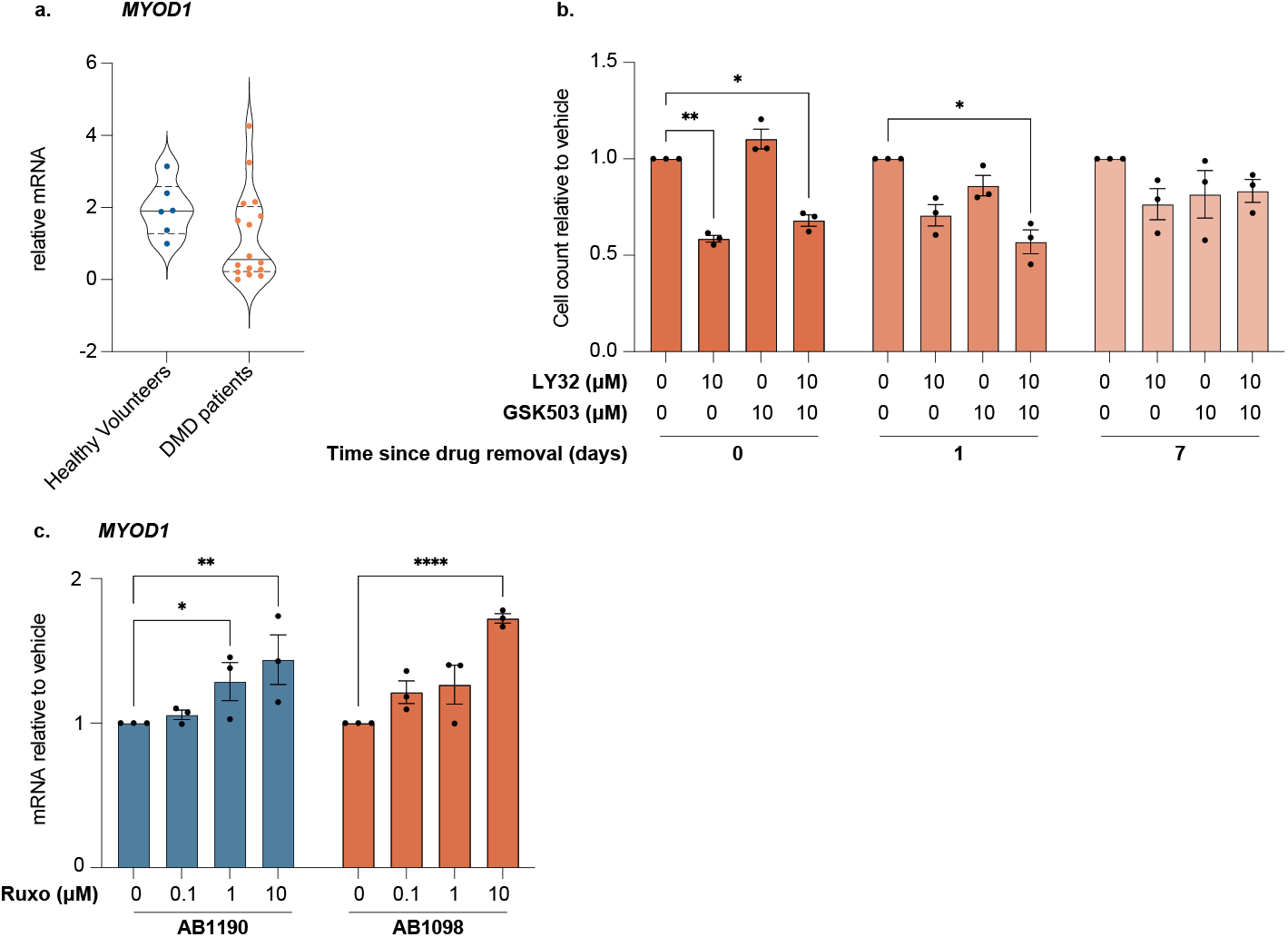
Ruxolitinib treatment causes increased *MYOD1* in proliferating myoblasts. **Corresponding to Figure 3. a**. Basal levels of *MYOD1* mRNA in primary healthy human myoblasts (n=6) and primary DMD patient-derived myoblasts (n=16). Lines show median and quartiles. **b**. AB1098 cells were treated with 10 μM LY32 +/- 10 μM GSK503 or vehicle (DMSO) for 24 hours. Directly following treatment (0 days since drug removal) cells were manually counted, with trypan blue staining to discount dead cells. Cells were then replated at a constant concentration, cultured in proliferation media without drugs for 24 hours and counted again (1 day since drug removal). Cells were then replated at a constant concentration and cultured again for 6 days before counting (7 days since drug removal). Cells were passaged once in this 6-day period. Cell count is shown relative to the vehicle only control for each time point. **c**. RT-qPCR of *MYOD1* in immortalised myoblasts AB1190 and AB1098 following treatment with varying concentrations of Ruxolitinib (Ruxo). Results are shown relative to vehicle treated sample with a two-way ANOVA with Dunnetts multiple comparison test to determine significance. Graph shows mean (n=3) +/- SEM. **a-b**. *MYOD1* is normalised against *TBP*. *p<0.05, ***p<0.001, ****p<0.0001.

**Supplementary Figure 4:**
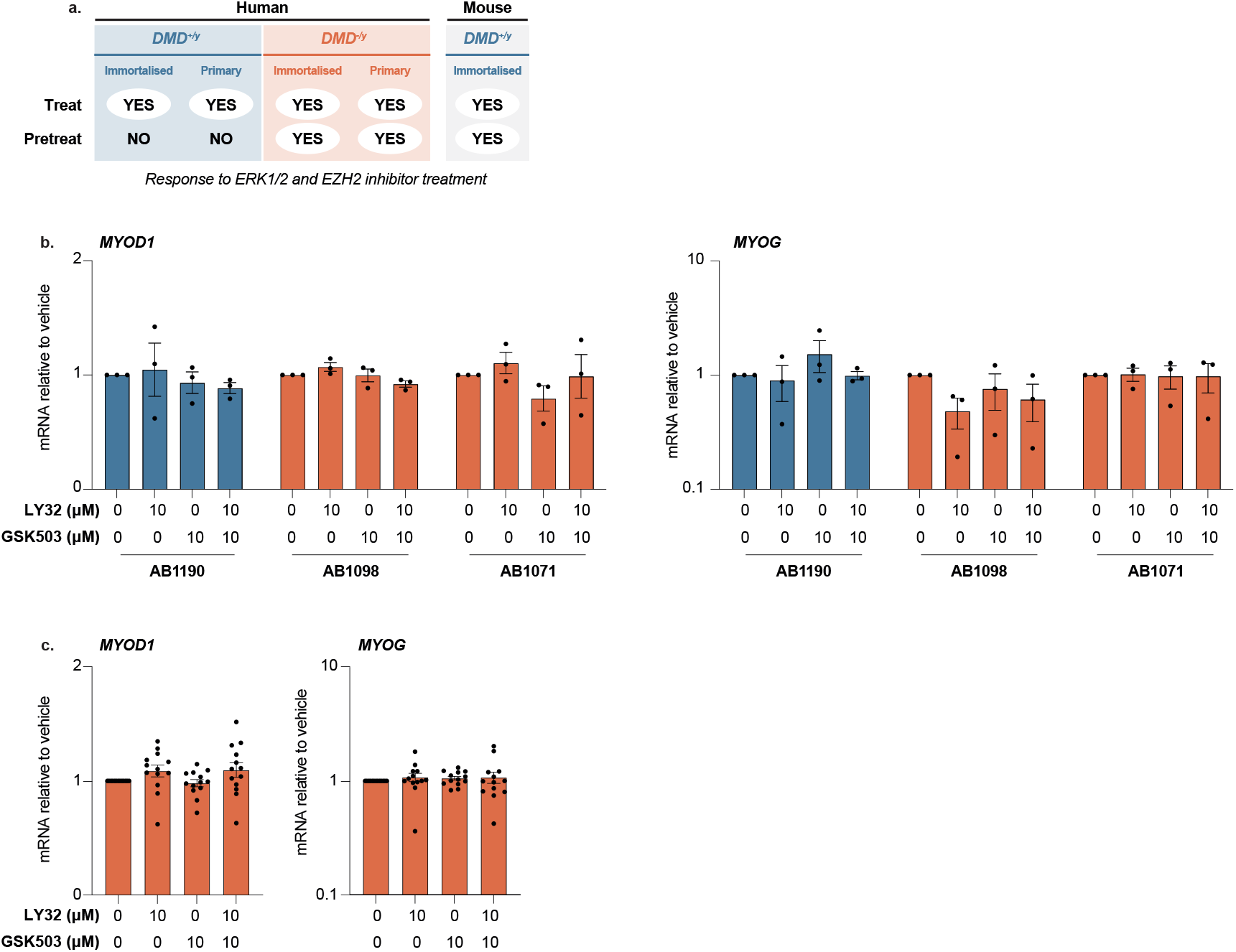
*MYOD1* and *MYOG* expression in differentiated myotubes are unaltered by pre-treatment with LY32 and GSK503. **a**. Table demonstrating the difference in sensitivities to LY32 +/- GSK503 treatment and pre-treatment between DMD myoblasts, healthy human myoblasts and healthy mouse myoblasts. Mouse results taken from ^23^. ‘YES’ indicates increase in *UTRN* to either or both drugs. **a-b**. Proliferating myoblasts were treated with LY32 +/- GSK503 or vehicle for 24 hours, following which drugs were removed and cells allowed to differentiate for 96 hours. Cells were sampled as myotubes following differentiation. RT-qPCR for *MYOD1* and *MYOG* mRNA following treatment. Results are normalised to *TBP* and shown relative to the vehicle pre-treated sample. Graphs show mean +/- SEM. **a**. Immortalised DMD (AB1098 and AB1071) and healthy control (AB1190) myoblasts (n=3). **b**. Primary DMD myoblasts (n=14).

## Methods

### Myoblast isolation

#### Primary human myoblasts from healthy human volunteers

Muscle samples were obtained from six young (25 ± 4 years) healthy male volunteers using the Bergstrom needle biopsy technique with additional suction from the vastus lateralis who gave written informed consent for their cells to be used in this study. All experiments were performed with UK NHS Ethics Committee approval (London Research Ethics Committee; reference: 16/LO/1707) and in accordance with the Human Tissue Act and Declaration of Helsinki.

Cell isolation was performed as in ^39^. The muscle biopsy was enzymatically digested in basal medium (C-23060; PromoCell, Heidelberg, Germany) containing collagenase D (2 mg/mL, Roche, Germany) and Dispase II (2 mg/mL, Sigma) for 1 h at 37°C with trituration every 15 min. After filtering through a 100-μm filter (BD Falcon) and centrifugation cells were resuspended in proliferation medium (15% FBS, C-23060; PromoCell, Heidelberg, Germany) and transferred to a T-25 tissue culture vessel (Nunc, Germany). Cells were maintained in a humidified incubator at 37°C and 6% CO_2_ for 7 days with media changes every 48 h, cells from the first change were collected by centrifugation, resuspended in fresh medium and returned to their original culture vessel.

At Day 7 post-biopsy, immuno-magnetic cell sorting (MACS) using CD56+ve Magnetic beads (Miltenyi Biotec) to separate CD56+ve (enriched for myoblasts) and CD56-ve (enriched for fibroblasts) fractions, as described previously.

#### Primary DMD patient-derived myoblasts

Primary DMD patient-derived myoblasts have been supplied by the MRC Centre for Neuromuscular Diseases (MRC CNMD) Biobank London (UCL, London, UK; Research Ethics Committee reference no.06/Q0406/33). For all samples collected by the Biobank after 01/09/2006, written consent for research has been obtained from all patients or their parent/guardian. All samples have been supplied to the project anonymised.

#### Immortalised myoblasts

Immortalised DMD patient-derived (AB1098DMD14S and AB1071DMD13PV, referred to as AB1098 and AB1071) and healthy control (AB1190) myoblasts were isolated from human biopsies provided by MyoBank, affiliated to EuroBioBank and authorized to distribute such material for research (Authorization AC-2019-3502 from the French Ministery for Higher Education, Research and innovation). Primary cultures were isolated and immortalized as already described ^40^.

### Myoblast culture

Proliferating myoblasts were cultured in skeletal muscle cell growth medium ready-to-use kit of basal medium plus supplement mix (Sigma Aldrich, C-23060), supplemented with 10% Fetal Bovine Serum (FBS). This was further supplemented with 0.5% penicillin-streptomycin (Gibco) and 0.2 mM L-glutamine (Gibco) for healthy human primary myoblasts and immortalised myoblasts. While primary DMD-derived myoblasts were cultured in complete skeletal muscle growth medium plus 10% FBS supplemented with 3 mM Glutamax I (Invitrogen) and 6 μg/ml Gentamicin (Sigma). All myoblasts were maintained in a 37 °C incubator with 5% CO_2_. Proliferating cultures were maintained at low confluency to prevent the onset of differentiation.

To promote differentiation, cells were grown to confluency in proliferation medium and then transferred to basal skeletal muscle cell growth medium without the addition of FBS or supplement mix. The relevant antibiotics (penicillin-streptomycin or gentamicin) were supplemented.

### Genotyping

DNA was extracted from cell pellets using DNeasy Blood and Tissue kit (Qiagen) as per the manufacturer’s instructions. PCR was performed using HotStart Taq Polymerase (Thermo Fisher Scientific) with a melting temperature of 58°C, using 50 ng of DNA and the primers pairs listed in Table 1.

**Table 1.**
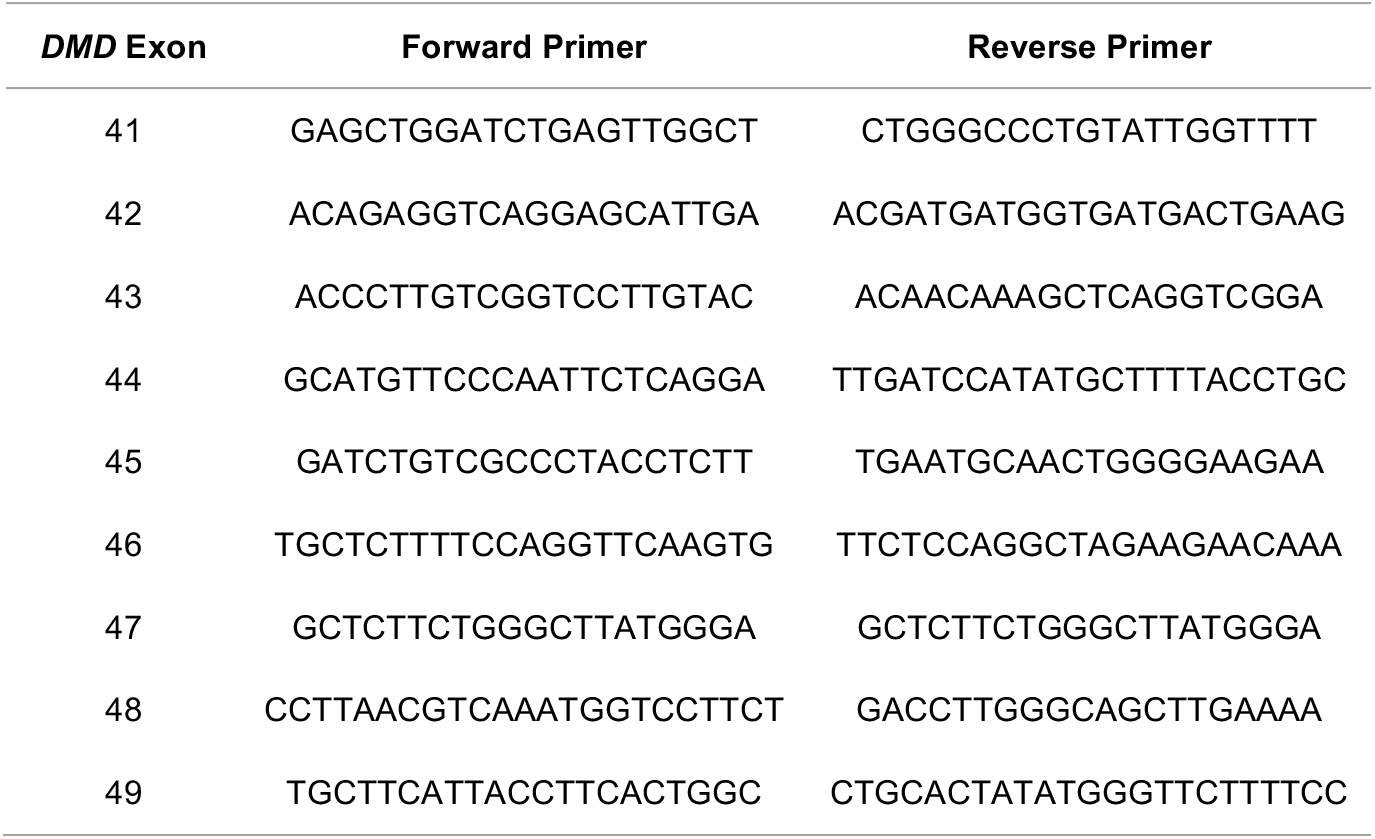
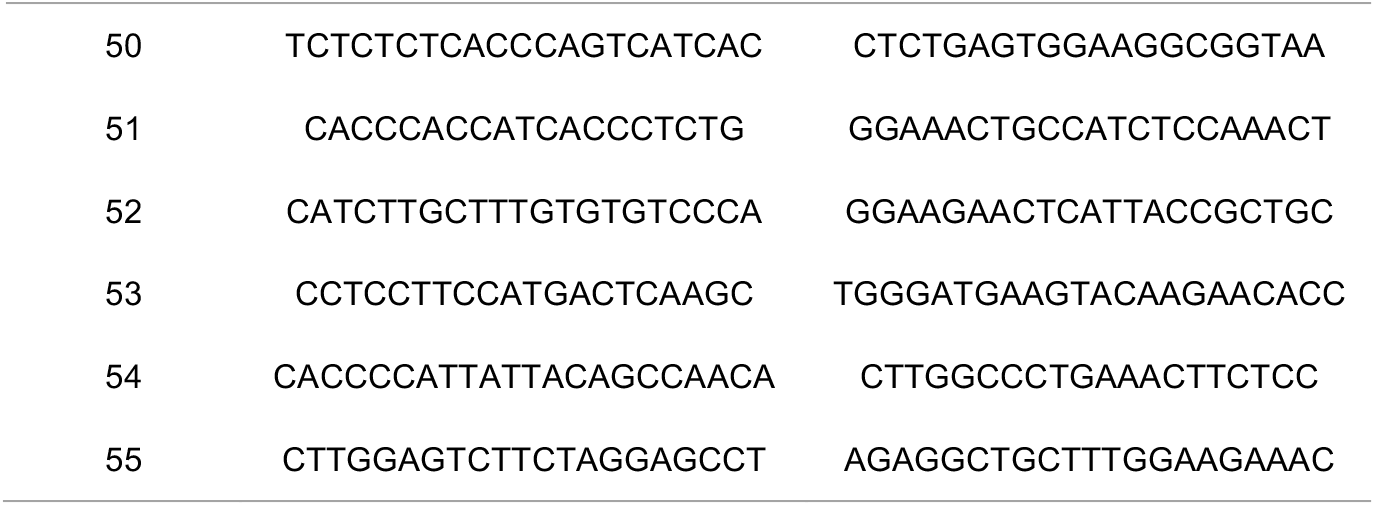
Primers used for *DMD* genotyping.

### Drug Treatments

LY3214996, GSK503 and Ruxolitinib were purchased from SelleckChem and resuspended in DMSO to generate 10 mM stock solutions. Proliferating myoblasts were plated and allowed to attach prior to treatment, then treated for 24 hours in proliferation medium. For pre-treatment experiments, drugs and proliferation media was then removed and replaced with differentiation media for 24-96 hours.

### Reverse transcription quantitative real-time PCR (RT-qPCR)

RNA was extracted using the RNeasy Mini Kit (Qiagen) with on column DNase digestion. Cells were lysed in 350 μl RLT buffer (Qiagen) and RNA extraction performed as the manufacturer described. SuperScript III reverse transcriptase kit, with 10 μM random primers (Thermo Fisher) was used to generate cDNA according to the manufacturer’s instructions. Next RT-qPCR was performed using the QuantiTect SYBR Green PCR mix (Qiagen) with 10 nM forward and reverse primers (Table 2). *TBP* was used as a house keeping gene and samples are shown relative to the vehicle-treated control using the formula EI^control Ct - sample Ct^/EH^control Ct - sample Ct^, where EI = efficiency of primers of interest and EH = efficiency of housekeeping primers. Efficiency was calculated using the formula E = 10^‐1/Slope^. All samples were run in triplicate.

**Table 2:**
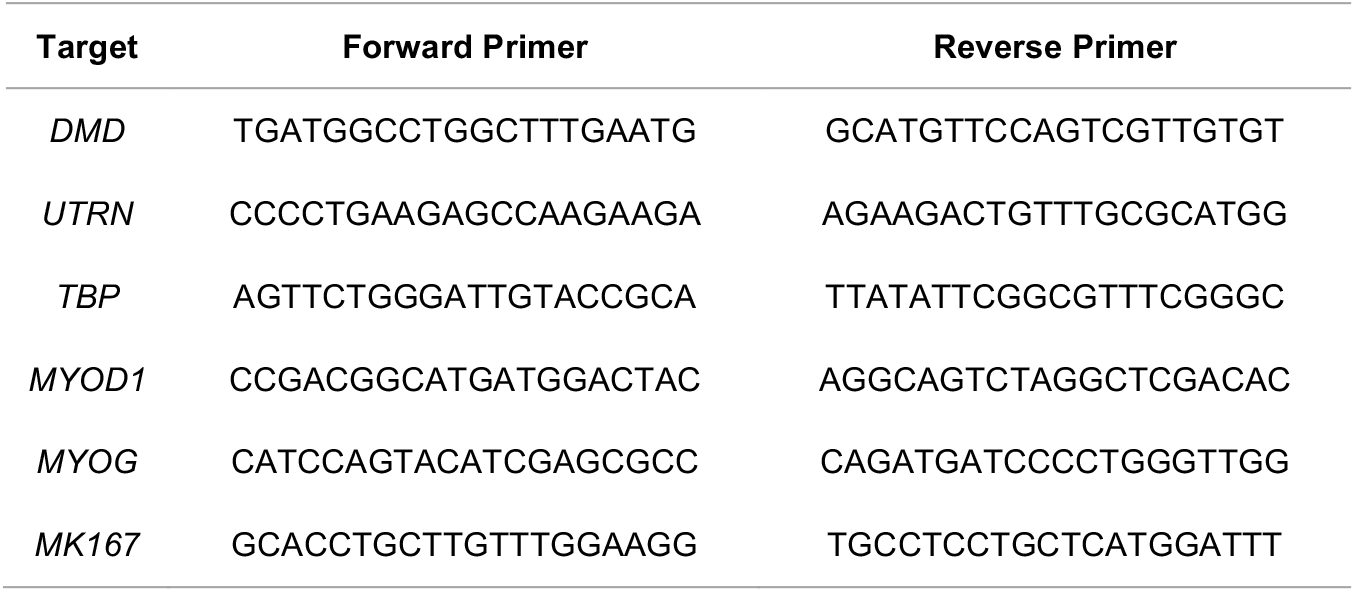
Primers used for RT-qPCR.

### Western Blot

Myoblasts were washed in cold PBS, then collected in cold PBS using a cell scraper. Cells were pelleted by centrifugation at 1200 rpm for 3 minutes and lysed in 150 μl RIPA buffer (Pierce, Thermo Scientific 89900) supplemented with protease and phosphatase inhibitors for 10 minutes at 4°C in rotation. Then samples were centrifuged at 15,000rpm at 4°C for 15 minutes. The protein concentration of the supernatant was determined using the BCA kit (Pierce Thermo Scientific 23225), as per the manufacturer’s instructions. Samples were diluted to 1.5-3 μg/μl in RIPA buffer plus 4X laemmli sample buffer (BioRad 1610747) and heated to 95°C for 5 minutes.

25-30 μg protein was loaded per sample onto a 4-20% polyacrylamide gel (BioRad) using a BioRad tank in 1X Tris/Glycine/SDS running buffer (BioRad). The gel was run for 30 minutes at 80 V followed by 2-3 h at 150 V. A precision plus dual colour (BioRad 1610374) and a HiMark (Invitrogen LC5699) ladder were run for each gel.

Prior to transfer each gel was cut to separate the dystrophin and utrophin bands (at 427 and 400 kDa, respectively) from the vinculin bands (at 116 kDa). The top part of the gel, containing dystrophin and utrophin was subject to wet transfer using BioRad apparatus. Transfer was run using 1X Tris/Glycine/SDS running buffer (BioRad) + 5% ethanol and gels were allowed to equilibrate in this at RT for 15 minutes prior to transfer. PVDF membrane (Millipore IPLL00010) was incubated in 100% ethanol for 5 minutes at RT. Transfer was run at 300 mA for 3 hours at 4°C and on ice. Dry transfer using the iBlot2 system and PVDF membranes (Invitrogen IB24002) was used for the lower part of the gel, containing vinculin. This was run at 20V for 7 minutes.

Both membranes were blocked in 3% BSA in TBST for 1 hour at RT. Membranes were then incubated in primary antibodies (Dystrophin: Cell signalling technologies, 47759, 1:500; Utrophin: Santa Cruz, Sc-33700 1:500; and Vinculin: Abcam, 129002, 1:10,000) in 3% BSA overnight at 4°C. The following day membranes were washed 3 × 7 minutes in TBST, then incubated with secondary antibodies (Goat anti-rabbit: IRDye 800CW, LICOR bio, 926-32211, 1:20,000, Goat anti-mouse: IRDye 680, LICOR bio, 926-68070, 1:10,000) in 3% BSA for 1 hour at RT, then washed again 3 × 7 minutes in TBST, followed by a final wash in dH_2_O. Bands were visualised using a LI-COR Odyssey Clx instrument with Image Studio 6 software. Gel densitometry was calculated using FIJI ^41^ (ImageJ2 version 2.14.0/1.54f).

### Immunofluorescent staining

Myoblasts were grown in standard culture conditions on glass-bottomed chamber slides (Nunc Lab-Tek) treated with 0.1mg/ml collagen (Corning) in 0.02N acetic acid prior to culture. Following culture, slides were washed twice in PBS, incubated with 4% formaldehyde for 10 minutes at RT, followed by a further three PBS washes. Cells were permeabilised in 1% BSA

+ 0.01% Trition X-100 for 15 minutes at RT, washed again 3X in PBS and blocked for 1 h in 1% BSA. Slides were incubated with primary antibodies overnight at RT in 1% BSA in PBS (MHC: DSHB MF20 1:200; Myogenin: DSHB, F5D, 1:50; Ki67: Abcam 15580 1:500). The following day, slides were washed 3X in PBS, and incubated with secondary antibodies (Goat anti-rabbit: AF568, Invitrogen A11011,1:500; Goat anti-mouse IgG2b: AF647, Invitrogen A21242, 1:500; Goat anti-mouse IgG1: AF488, Invitrogen A21131, 1:500) for 1 hr at RT. Antibodies were removed with a further 3 washes in PBS and slides were incubated with DAPI for 10 minutes at RT. Slides were washed a further 2X in PBS prior to mounting with Vectashield (Vector Labs H-1000).

Images were taken using an SP5 confocal microscope (Leica) and analysed using FIJI ^41^ (ImageJ2 version 2.14.0/1.54f). For Ki67 nuclear intensity, all images were first converted to grey scale. A duplicate copy of the DAPI image was created and converted to binary to identify nuclei. For quantification of Ki67 and DAPI signal, the analyse particle function was used: the binary DAPI image was redirected to the grey scale Ki67 or DAPI image and integrated density calculated for each nucleus. A minimum of 3 images were analysed for each sample and all individual nuclei are represented in graphs.

### Cell count and proliferation

AB1098 cells were plated in 12 well plates and treated with 10 μM LY32 +/- 10 μM GSK503 or vehicle (DMSO) for 24 hours. Following treatment, media and drugs were removed and cells were resuspended in fresh media. An aliquot was mixed 1:1 in trypan blue and cell number manually counted using a haemocytometer. For each condition 5 × 10^4^ cells were replated in a well of a 12 well plate and cultured in proliferation media for 24 hours. The following day cells were counted again, replated at 1 × 10^4^ cells per well and cultured in proliferation media for 3 days. Cells were replated at 1 × 10^4^ cells per well and cultured for a further 3 days. At day 7 post drug removal, cells were collected and counted a final time.

### Statistics and reproducibility

Microsoft Excel was used for calculations of mean and relative values. GraphPad Prism (Version 10.2.2) was used to generate graphs and perform statistical analysis. Experiments involving immortalised myoblasts were performed in triplicate and graphs show the mean for each cell line. For primary myoblasts, an individual experiment was run for each biopsy and graphs present the mean for all healthy (n=6) or DMD patient-derived (n=16) biopsies to show population-based trends. One or two-way ANOVAs were used for multi-condition comparisons (for example response to LY32 +/- GSK503 treatment) and unpaired T-tests were used when only two means were compared (for example between cell lines or genotypes). The specific details of each test used are outlined in the figure legends.

## Notes

### Competing Interest Statement

The authors have declared no competing interest.

## References

1 Allikian, M. J. & McNally, E. M. Processing and assembly of the dystrophin glycoprotein complex. Traffic 8, 177–183 (2007).10.1111/j.1600-0854.2006.00519.x

2 Morgan, J. E. & Zammit, P. S. Direct effects of the pathogenic mutation on satellite cell function in muscular dystrophy. Exp Cell Res 316, 3100–3108 (2010).10.1016/j.yexcr.2010.05.014

3 Gao, Q. Q. & McNally, E. M. The Dystrophin Complex: Structure, Function, and Implications for Therapy. Compr Physiol 5, 1223–1239 (2015).10.1002/cphy.c140048

4 Hoffman, E. P., Brown, R. H., Jr. & Kunkel, L. M. Dystrophin: the protein product of the Duchenne muscular dystrophy locus. Cell 51, 919–928 (1987).10.1016/0092-8674(87)90579-4

5 Duan, D., Goemans, N., Takeda, S., Mercuri, E. & Aartsma-Rus, A. Duchenne muscular dystrophy. Nat Rev Dis Primers 7, 13 (2021).10.1038/s41572-021-00248-3

6 Swiderski, K. & Lynch, G. S. Murine models of Duchenne muscular dystrophy: is there a best model? Am J Physiol Cell Physiol 321, C409–C412 (2021).10.1152/ajpcell.00212.2021

7 Yucel, N., Chang, A. C., Day, J. W., Rosenthal, N. & Blau, H. M. Humanizing the mdx mouse model of DMD: the long and the short of it. NPJ Regen Med 3, 4 (2018).10.1038/s41536-018-0045-4

8 Sicinski, P. et al. The molecular basis of muscular dystrophy in the mdx mouse: a point mutation. Science 244, 1578–1580 (1989).10.1126/science.2662404

9 Anderson, J. E., Bressler, B. H. & Ovalle, W. K. Functional regeneration in the hindlimb skeletal muscle of the mdx mouse. J Muscle Res Cell Motil 9, 499–515 (1988).10.1007/BF01738755

10 Deconinck, A. E. et al. Utrophin-dystrophin-deficient mice as a model for Duchenne muscular dystrophy. Cell 90, 717–727 (1997).10.1016/s0092-8674(00)80532-2

11 Clerk, A., Morris, G. E., Dubowitz, V., Davies, K. E. & Sewry, C. A. Dystrophin-related protein, utrophin, in normal and dystrophic human fetal skeletal muscle. Histochem J 25, 554–561 (1993).

12 Pons, F., Nicholson, L. V., Robert, A., Voit, T. & Leger, J. J. Dystrophin and dystrophin-related protein (utrophin) distribution in normal and dystrophin-deficient skeletal muscles. Neuromuscul Disord 3, 507–514 (1993).10.1016/0960-8966(93)90106-t

13 Ohlendieck, K. et al. Dystrophin-related protein is localized to neuromuscular junctions of adult skeletal muscle. Neuron 7, 499–508 (1991).10.1016/0896-6273(91)90301-f

14 Song, Y. et al. Non-immunogenic utrophin gene therapy for the treatment of muscular dystrophy animal models. Nat Med 25, 1505–1511 (2019).10.1038/s41591-019-0594-0

15 Deconinck, N. et al. Expression of truncated utrophin leads to major functional improvements in dystrophin-deficient muscles of mice. Nat Med 3, 1216–1221 (1997).10.1038/nm1197-1216

16 Fisher, R. et al. Non-toxic ubiquitous over-expression of utrophin in the mdx mouse. Neuromuscul Disord 11, 713–721 (2001).10.1016/s0960-8966(01)00220-6

17 Tinsley, J. et al. Expression of full-length utrophin prevents muscular dystrophy in mdx mice. Nat Med 4, 1441–1444 (1998).10.1038/4033

18 Cerletti, M. et al. Dystrophic phenotype of canine X-linked muscular dystrophy is mitigated by adenovirus-mediated utrophin gene transfer. Gene Ther 10, 750–757 (2003).10.1038/sj.gt.3301941

19 Tinsley, J. M. et al. Daily treatment with SMTC1100, a novel small molecule utrophin upregulator, dramatically reduces the dystrophic symptoms in the mdx mouse. PLoS One 6, e19189 (2011).10.1371/journal.pone.0019189

20 Muntoni, F. et al. PhaseOut DMD: a Phase 2, proof of concept, clinical study of utrophin modulation with ezutromid. Neuromuscular Disord 27, S217–S217 (2017).10.1016/j.nmd.2017.06.443

21 Perkins, K. J. & Davies, K. E. Alternative utrophin mRNAs contribute to phenotypic differences between dystrophin-deficient mice and Duchenne muscular dystrophy. FEBS Lett 592, 1856–1869 (2018).10.1002/1873-3468.13099

22 Guiraud, S. & Davies, K. E. Pharmacological advances for treatment in Duchenne muscular dystrophy. Curr Opin Pharmacol 34, 36–48 (2017).10.1016/j.coph.2017.04.002

23 Gleneadie, H. J. et al. Endogenous bioluminescent reporters reveal a sustained increase in utrophin gene expression upon EZH2 and ERK1/2 inhibition. Commun Biol 6 (2023).ARTN 318 10.1038/s42003-023-04666-9

24 Stern-Straeter, J., Bonaterra, G. A., Hormann, K., Kinscherf, R. & Goessler, U. R. Identification of valid reference genes during the differentiation of human myoblasts. BMC Mol Biol 10, 66 (2009).10.1186/1471-2199-10-66

25 Eigler, T. et al. ERK1/2 inhibition promotes robust myotube growth via CaMKII activation resulting in myoblast-to-myotube fusion. Dev Cell 56, 3349–3363 e3346 (2021).10.1016/j.devcel.2021.11.022

26 Dumont, N. A. et al. Dystrophin expression in muscle stem cells regulates their polarity and asymmetric division. Nature Medicine 21, 1455-+ (2015).10.1038/nm.3990

27 Guiraud, S. et al. The Pathogenesis and Therapy of Muscular Dystrophies. Annu Rev Genomics Hum Genet 16, 281–308 (2015).10.1146/annurev-genom-090314-025003

28 Gosselin, M. R. F. et al. Loss of full-length dystrophin expression results in major cell-autonomous abnormalities in proliferating myoblasts. Elife 11 (2022).10.7554/eLife.75521

29 Sun, L. et al. JAK1-STAT1-STAT3, a key pathway promoting proliferation and preventing premature differentiation of myoblasts. J Cell Biol 179, 129–138 (2007).10.1083/jcb.200703184

30 Falcucci, L. et al. Transcriptional adaptation upregulates utrophin in Duchenne muscular dystrophy. Nature 639, 493–502 (2025).10.1038/s41586-024-08539-x

31 Helliwell, T. R., Man, N. T., Morris, G. E. & Davies, K. E. The dystrophin-related protein, utrophin, is expressed on the sarcolemma of regenerating human skeletal muscle fibres in dystrophies and inflammatory myopathies. Neuromuscul Disord 2, 177–184 (1992).10.1016/0960-8966(92)90004-p

32 Guiraud, S., Roblin, D. & Kay, D. E. The potential of utrophin modulators for the treatment of Duchenne muscular dystrophy. Expert Opin Orphan D 6, 179–192 (2018).10.1080/21678707.2018.1438261

33 Schuierer, M. M., Mann, C. J., Bildsoe, H., Huxley, C. & Hughes, S. M. Analyses of the differentiation potential of satellite cells from myoD-/-,mdx, and PMP22 C22 mice. BMC Musculoskelet Disord 6, 15 (2005).10.1186/1471-2474-6-15

34 Megeney, L. A., Kablar, B., Garrett, K., Anderson, J. E. & Rudnicki, M. A. MyoD is required for myogenic stem cell function in adult skeletal muscle. Genes Dev 10, 1173–1183 (1996).10.1101/gad.10.10.1173

35 Megeney, L. A. et al. Severe cardiomyopathy in mice lacking dystrophin and MyoD. Proc Natl Acad Sci U S A 96, 220–225 (1999).10.1073/pnas.96.1.220

36 Kodippili, K. & Rudnicki, M. A. Satellite cell contribution to disease pathology in Duchenne muscular dystrophy. Front Physiol 14 (2023).ARTN 1180980 10.3389/fphys.2023.1180980

37 Ghahramani Seno, M. M. et al. RNAi-mediated knockdown of dystrophin expression in adult mice does not lead to overt muscular dystrophy pathology. Hum Mol Genet 17, 2622–2632 (2008).10.1093/hmg/ddn162

38 Francis, T., Soendenbroe, C., Lazarus, N. R., Mackey, A. L. & Harridge, S. D. R. Insights into human muscle biology from human primary skeletal muscle cell culture. Journal of Muscle Research and Cell Motility 46, 301–318 (2025).10.1007/s10974-025-09696-w

39 Francis, T. G., Jaka, O., Ellison-Hughes, G. M., Lazarus, N. R. & Harridge, S. D. R. Human primary skeletal muscle-derived myoblasts and fibroblasts reveal different senescent phenotypes. JCSM Rapid Communications 5, 226–238 (2022).10.1002/rco2.67

40 Mamchaoui, K. et al. Immortalized pathological human myoblasts: towards a universal tool for the study of neuromuscular disorders. Skelet Muscle 1 (2011).10.1186/2044-5040-1-34

41 Schindelin, J. et al. Fiji: an open-source platform for biological-image analysis. Nat Methods 9, 676–682 (2012).10.1038/Nmeth.2019

